# Common coupling map advances GPCR-G protein selectivity

**DOI:** 10.1101/2021.09.07.459250

**Authors:** Alexander S. Hauser, Charlotte Avet, Claire Normand, Arturo Mancini, Asuka Inoue, Michel Bouvier, David E. Gloriam

## Abstract

Two-thirds of human hormones and one-third of clinical drugs act on membrane receptors that couple to G proteins to achieve appropriate functional responses. While G protein transducers from literature are annotated in the Guide to Pharmacology database, two recent large-scale datasets now expand the receptor-G protein ‘couplome’. However, these three datasets differ in scope and reported G protein couplings giving different coverage and conclusions on GPCR-G protein signaling. Here, we report a common coupling map uncovering novel couplings supported by both large-scale studies, the selectivity/promiscuity of GPCRs and G proteins, and how the co-coupling and co-expression of G proteins compare to the families from phylogenetic relationships. The coupling map and insights on GPCR-G protein selectivity will catalyze advances in receptor research and cellular signaling towards the exploitation of G protein signaling bias in design of safer drugs.

## Introduction

G protein-coupled receptors (GPCRs) represent the largest family of proteins involved in signal propagation across biological membranes. They recognize a vast diversity of signals going from photons and odors to neurotransmitters, hormones, and cytokines (1). Their main signaling modality involves the engagement and activation of G proteins. G proteins are heterotrimeric proteins consisting of a α, β and γ subunits that dissociate to α and a βγ upon activation by a GPCR. G proteins are named by their α subunit (16 in human) and are divided into four families which share homology and downstream signaling pathways: G_s_ (G_s_ and G_olf_), G_i/o_ (G_i1_, G_i2_, G_i3_, G_o_, G_z_, G_t1_, G_t2_ and G_gust_), G_q/11_ (G_q_, G_11_, G_14_ and G_15_) and G_12/13_ (G_12_ and G_13_). A GPCR’s activation of G proteins can be very selective or promiscuous and change upon ligand-dependent biased signaling that alters its profile on the G protein subtype or family levels. The pleiotropic signaling and ligand-dependent bias of GPCRs pose a grand challenge in human biology to map the differential activation of specific G proteins.

The IUPHAR/BPS Guide to Pharmacology (GtP) database contains reference data from expert curation of literature (1). GtP couplings covers 253 GPCRs and the four G protein families. The G protein families have been classified into “primary” and “secondary” transducers without quantitative values. Recently, the Inoue group determined the first large-scale quantitative coupling profiles of 148 GPCR_s_ to the G_q_ wildtype and 10 G protein chimeras employing a TGF-α shedding assay (2) (NTS_1_ and TRH_1_ added herein making it 150 receptors). Those chimeras consist of a G_q_ backbone in which the six most C-terminal Gα residues – a part of the H5 domain inserting to the intracellular receptor cavity – have been replaced to represent all 16 human G proteins (five of which have identical sequences to other G proteins, see below). In a paper accompanying the present analysis, the Bouvier group quantified the couplings of 100 GPCRs to 12 G proteins: G_s_, G_i1_, G_i2_, G_o_ (G_oA_ and G_oB_ isoforms), G_z_, G_q_, G_11_, G_14_, G_15_, G_12_ and G_13_ but not G_olf_ (couples mainly to olfactory receptors), G_i3_ and G_t1-2_ (Transducin, couples to rhodopsin (visual) receptors) and G_gust_ (Gustducin, couples to taste receptors) (3). The authors used novel enhanced bystander bioluminescence resonance energy transfer biosensors that allow to monitor G protein activation (G protein Effector Membrane Translocation assay; GEMTA) without need to modify the G protein subunits (except for G_s_) or the receptors.

Here, we analyze the GtP, Inoue and Bouvier coupling datasets to determine confident couplings supported by at least two independent sources, including novel couplings discovered jointly by the two latter sources. We establish a scalable protocol to normalize quantitative G protein couplings, combine E_max_ and EC_50_ into a common log(E_max_/EC_50_) (4) value and aggregate subtypes to allow comparisons across G protein families. On this basis, we develop a unified map of GPCR-G protein couplings that can be filtered or intersected in GproteinDb (5), describe GPCR-G protein selectivity across an unprecedented number of receptors and coupling data points and reveal correlated co-couplings.

## Results

### Tools and Resources – Common coupling map unifying GPCR-G protein data Systematic profiling doubles the average number of G protein family couplings of GPCRs

To obtain an overview of the coverages of GPCR-G protein coupling sources, we compared all couplings reported by both the Bouvier (3) and Inoue (2) groups and annotated in the Guide to Pharmacology database (GtP (1)) (Fig. 1). This shows that the three sources together comprise couplings for 265 (67%) out of the 398 non-olfactory GPCRs and that 70 of these receptors are present in all datasets (Fig. 1B). The Bouvier and Inoue datasets have collectively quantified individual G protein couplings of 178 receptors using one assay, whereas the remaining 87 receptors have so far only been annotated in GtP on the G protein family level from a multitude of publications and assays.

**Fig. 1.**
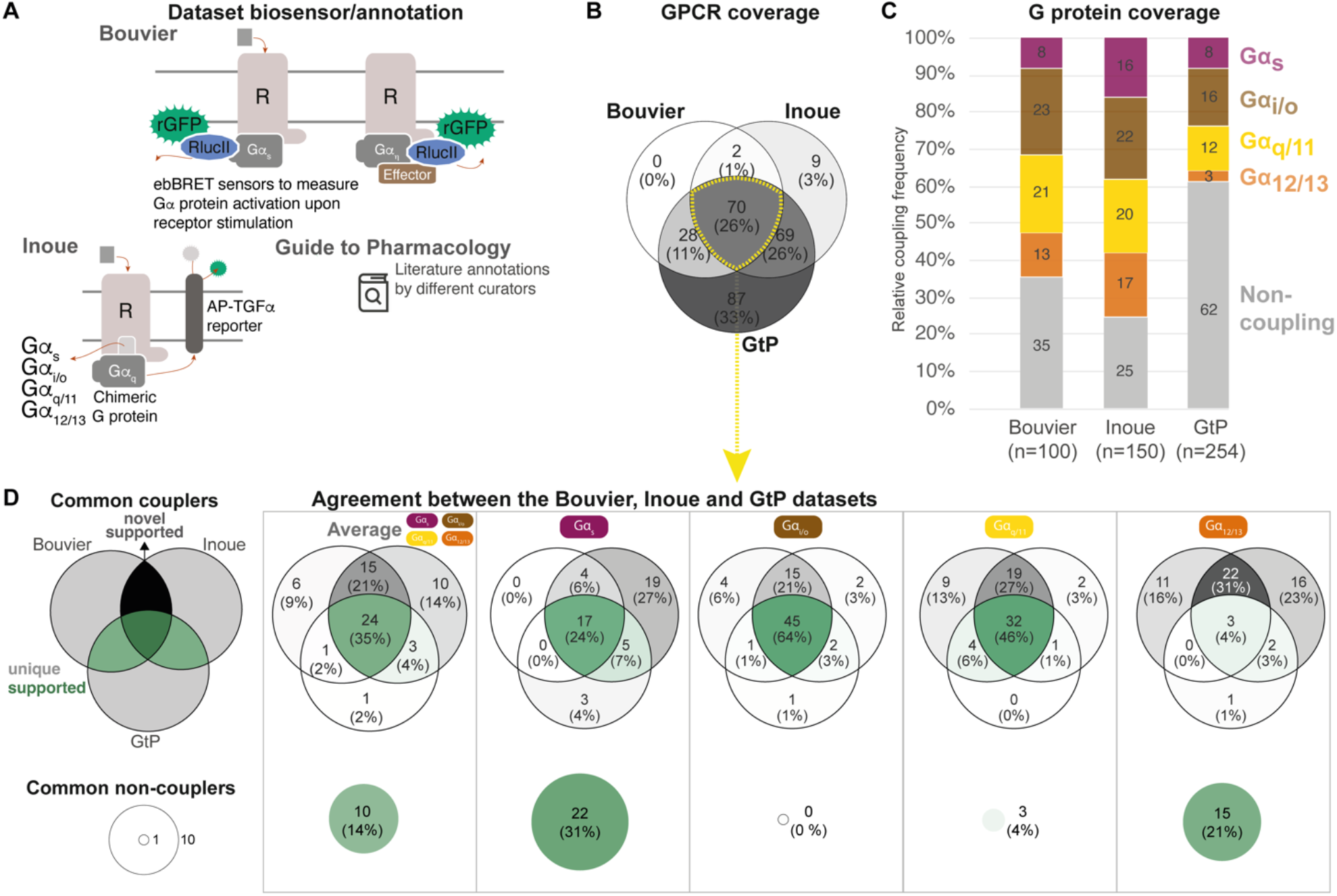
Coverage and agreement of the Bouvier lab, Inoue group and Guide to Pharmacology datasets. (**A**) Biosensor principles used by the Bouvier and Inoue groups, and literature annotation stored in the Guide to Pharmacology database (GtP). (**B**) Intersection of the GPCRs included in the Bouvier (n=100), Inoue (n=150) and GtP (n=254) datasets. (**C**) Relative family distributions of G protein couplings across datasets. **(D)** Comparison of the G protein family coupling profiles of 70 GPCRs present in all of the Bouvier, Inoue and GtP datasets. More detailed analysis of common G protein couplings for just the Bouvier and Inoue datasets are given in Fig. S1. **(A-D)** Note: All analyses herein cover the 12 G proteins: G_s_, G_i1_, G_i2_, G_oA_, G_oB_, G_z_, G_q_, G_11_, G_14_, G_15_, G_12_ and G_13_. G_olf_ and G_i3_ could not be analyzed, as they had not been tested by the Bouvier group. The Inoue data for the pairs G_i1_-G_i2_, G_oA_-G_oB_ and G_q_-G_11_ were generated with identical chimera inserting the G C-terminal hexamer into a G_q_ backbone (2) meaning that identical datapoints were used to assess their coverage and agreement with other datasets (Table S1).

To allow the comparison of coupling densities and distributions across datasets, we selected the E_max_ threshold (1.4 standard deviations above basal) that gives the best agreement between the Bouvier and Inoue dataset. We believe that this is the best possible means to estimate what is correct data (rather than false negative/positive couplings), as large-scale information about what G proteins and GPCRs couple physiologically is not available. This cut-off is more stringent than the minimum of 3% signal above basal used in Inoue’s original report (2). We also aggregated G protein subtype couplings of the families (see Methods). This reveals that while GtP covers the largest number of receptors they have relatively few couplings – only 38% of all GPCR-G protein family pairs are ‘couplers’ compared to 65% in the Bouvier and 75% in the Inoue dataset (average of 1.5, 2.6 and 3.0 G protein families per GPCR, respectively; Fig. 1C). In particular G_12/13_ couplings are underrepresented in GtP where they account for 3% of GPCR-G protein pair datapoints compared to 13% in Bouvier and 17% in Inoue. The Bouvier and Inoue datasets share a more similar overall distribution of couplings, expect for G_s_ coupling which is twice as frequent within the latter dataset (16% compared to 8%). This demonstrates that the two first systematic coupling profiling studies have substantially expanded the known GPCR-G protein ‘couplome’ and that their assay platforms are amenable to high-throughput profiling also for the G_12/13_ family for which robust activation assays appeared only recently (6–8).

Given that the Bouvier and Inoue groups used different biosensors and GtP annotates literature reports from very diverse assays, we sought to determine to which extent they report the same couplings for the same GPCRs i.e., their 70 common receptors (data in tab ‘BIG-QualComp’ in Spreadsheet S3). We find that all three sources agree on an average of 49% of G protein family couplings/non-couplings (distributed as G_s_: 55%, G_i/o_: 64%, G_q/11_: 50% and G_12/13_: 25%, Fig. 1D). When instead analyzing the quantitative studies alone (excluding GtP), the agreement increases to 70% across G protein families. This agreement is 68% for individual G protein datapoints, which display an even larger span ranging from at least 53% for G_15_ up to 81% for G_oA_ (Fig. S1). These findings define a sizeable reference set of consensus G protein couplings and show that consistent large-scale profiling studies generate more comparable results than literature.

### Common coupling map unifies GPCR-G protein datasets and opens analysis

To enable quantitative correlation of the Bouvier, Inoue and future couplings, we established a data processing protocol giving the highest similarity and correlation of coupling measurements across datasets (Methods). This uses log transformed EC_50_ values and minimum-maximum normalized E_max_ values combined into a unified log(E_max_/EC_50_) value and an aggregation of G proteins onto families based on the maximum subtype values (Fig. 2A). Based on this protocol, we created a common coupling map, integrating the Bouvier, Inoue and GtP datasets (Fig. 2B). This unified coupling map establishes that it is possible to obtain comparable quantitative values despite the differences between biosensors and enables quantitative cross-study comparisons herein and in future studies from the field.

**Fig. 2.**
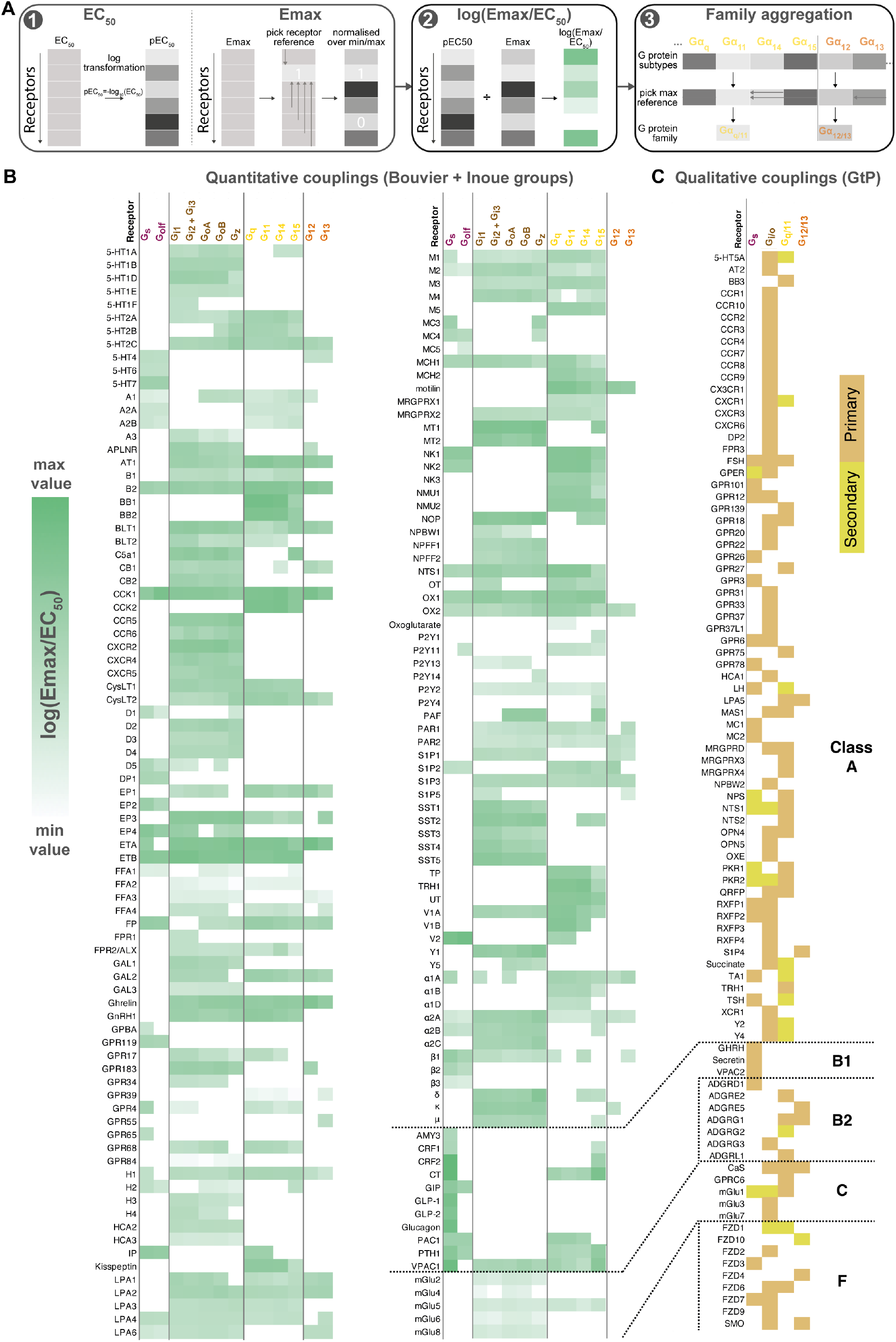
Map of GPCR-G protein couplings supported by at least two studies. (**A**) Normalization approach. E_max_ minimum-maximum normalization, E_C50_ log transformation and use of log(E_max_/E_C50_) as a combined measure with member-to-family aggregation by maximum G protein value. (**B**) Heatmap representation of log(E_max_/E_C50_) values for 166 GPCRs tested by the Bouvier and/or Inoue labs (for couplings with dual data sources a mean is used). For GtP, a G protein subtype is considered supported if the family has a known coupling. G_i2_ and G_i3_ are represented by the same / identical chimeric G protein in Inoue’s dataset (Table S1). (**C**) Heatmap representation of primary and secondary transducers for 90 GPCRs which couplings are only covered by the GtP database. (**B-C**) Empty cells (white) indicate no coupling. All source values are available in tab ‘Fig_4’ in Spreadsheet S5. Note: Researchers wishing to use this coupling map, optionally after applying own reliability criteria or cut-offs, can do so for any set of couplings in GproteinDb (5).

To enable any researcher to use the coupling map, we have availed a “G protein couplings” browser (https://gproteindb.org/signprot/couplings) in GproteinDb (7). By default, the coupling map (Fig. 2B) and browser only shows “supported” couplings with evidence from two datasets, but there is an option (first blue button) to changes the level of support to only one (for most complete coverage of GPCRs) or to three (for the highest confidence) sources. An exception, GtP couplings do not require additional support, as they are in most cases supported by multiple independent publications. We propose a standardized terminology to describe couplings based on their level of experimental support from independent groups (Table 1). The criterion of supporting independent data, and the terms “proposed” and “supported”, are already used by the Nomenclature Committee of the International Union of Basic and Clinical Pharmacology (NC-IUPHAR) for GPCR deorphanization. Furthermore, the online coupling browser allows any researcher to use only a subset of datasets, or to apply filters to the Log(Emax/EC50), Emax, and EC50 values. Finally, users can filter datapoints based on a statistical reliability score in the form of the number of SDs from basal response.

**Table 1:** Terms and definitions used to classify GPCR-G protein couplings.

| **Term** | **Definition** |
| --- | --- |
| Supported | Coupling or non-coupling datapoints that is supported by at least one other dataset i.e., at least 2 in total. An exception is made for couplings reported only in the GtP database, and not yet tested in a quantitative dataset, as the couplings in GtP are in most cases supported by multiple independent publications. |
| Novel | Coupling that is supported by Bouvier and Inoue but not present in GtP. |
| Proposed | Coupling identified in one but not yet tested in a second quantitative dataset (here Bouvier or Inoue). This refers to mainly receptors, but also G protein subtypes, that have only been tested in one quantitative dataset. Couplings reported only in the GtP database are not considered “proposed” but “supported” because they in most cases are based on the annotation of multiple independent publications. |
| Unique | Coupling that is unique in one dataset (other datasets have no coupling). |
| Missing | Non-coupling in the given dataset but coupling in the other compared datasets. |

## Research Advances – Insights on GPCR-G protein selectivity

### The Bouvier and Inoue datasets jointly support 101 novel G protein couplings

We next identified the ‘novel’ G protein couplings for which a family annotation is missing in GtP but have high confidence from dual support by the Bouvier and Inoue groups (Fig. S1). This revealed 38 receptors with novel couplings to 101 G proteins distributed across all families: G_s_,: 4, G_i/o_: 15, G_q/11_: 10 and G_12/13_: 21 (Fig. 3). The largest expansions – an increase by three of G protein families – was obtained for the histamine H_1_ and endothelin ET_A_ receptors which was found to couple to all G protein families but only have G_q/11_-coupling in GtP. Whereas it could be expected that GtP would miss couplings, we also analyzed if its expert curation excluded couplings that may be false positives as they are contradicted by both quantitative studies. This uncovered such G_s_-coupling to the α_2C_-adrenoceptor and cannabinoid CB_1-2_ receptors, G_12/13_ coupling to the purinergic P2Y_2_ receptor and G_i/o_-coupling to the β_2_-adrenoceptor, which however had weak G_z_ and G_oB_ coupling in the Bouvier study but did not cross the signal threshold (3). Notably, this is only 2% (5/254) of all GtP’s GPCR-G protein family pairs. Taken together, these findings serve to quantify the expansion of the GPCR-G protein ‘couplome’ while also confirming the outstanding accuracy of the expert annotation in the GtP database. Therefore, the large number of new couplings are mainly due to missing investigations in literature rather than incorrect curation.

**Fig. 3.**
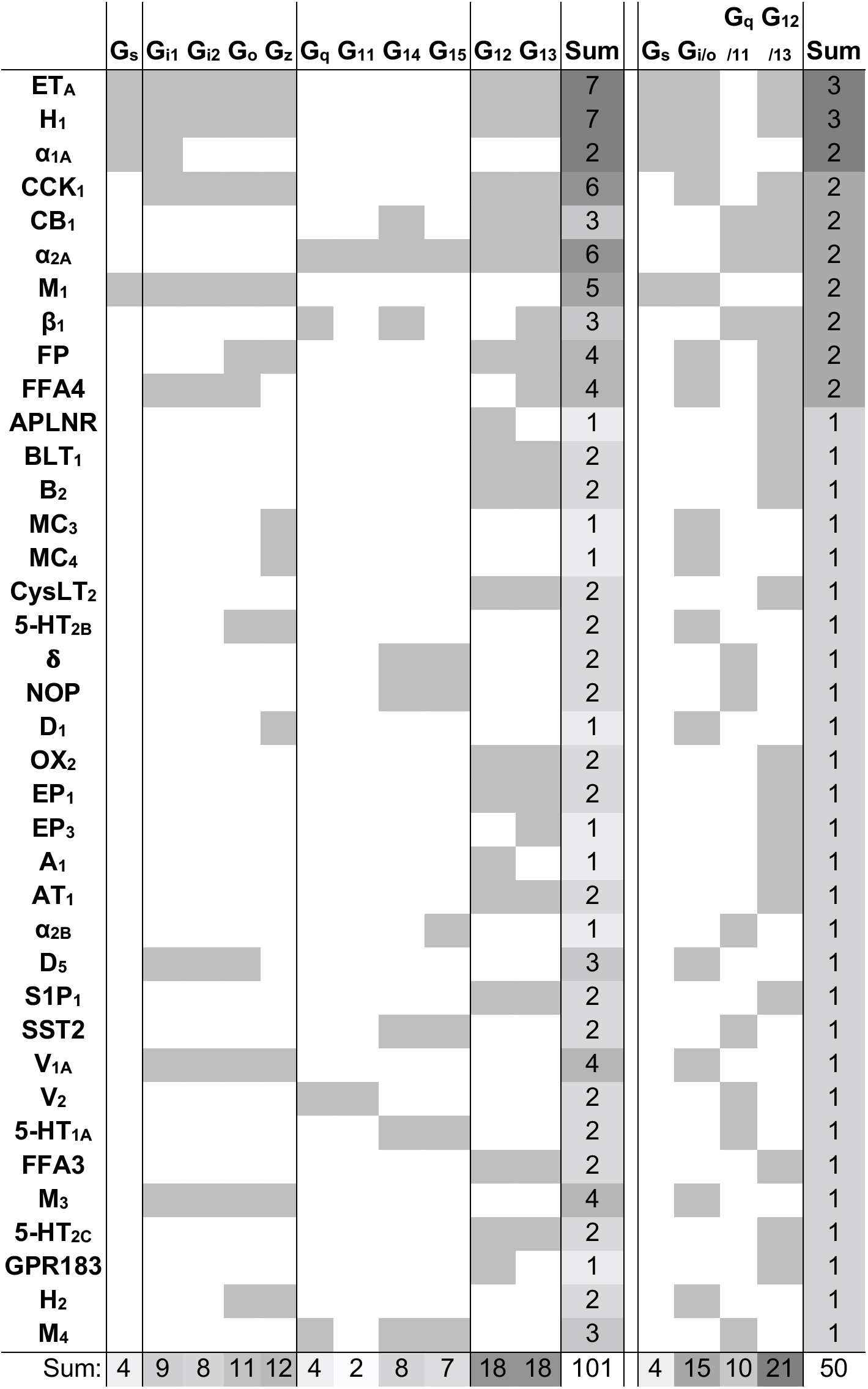
Novel G protein couplings identified by both the Bouvier and Inoue groups. Novel couplings identified by the Bouvier and Inoue groups but not in Guide to Pharmacology. G protein families are here considered shared if at least one specific subtype is found to couple in both dataset, which is not the case for the bottommost four receptors.

### Half of GPCRs are selective for a single whereas 5% promiscuously activate all G protein families

To gain insight into their levels of coupling selectivity, we intersected the G protein profiles of all receptors and counted the number of coupling partners for GPCRs and G proteins (Fig. 4). On the G protein family level (topmost in Fig. 4), our analysis spans 90 receptors with data only in GtP and 166 GPCRs with data from the Bouvier and/or Inoue groups – totaling 256 receptors. We require couplings from the Bouvier and Inoue groups to be supported by a second dataset (GtP couplings are already typically supported by several publications). We find that these 256 GPCRs couple to on average 1.7 G protein families distributed as 126 single-, 83 double-, 34 triple- and 13 all-family activating receptors. The share of fully selective (single-family activating) receptors differs largely across G protein families spanning from 6% for G_12/13_ to 22% for G_q/11_, 26% for G_s_ and up to 40% for G_i/o_ (tab ‘Fig_4’ in Spreadsheet S5).

**Fig. 4.**
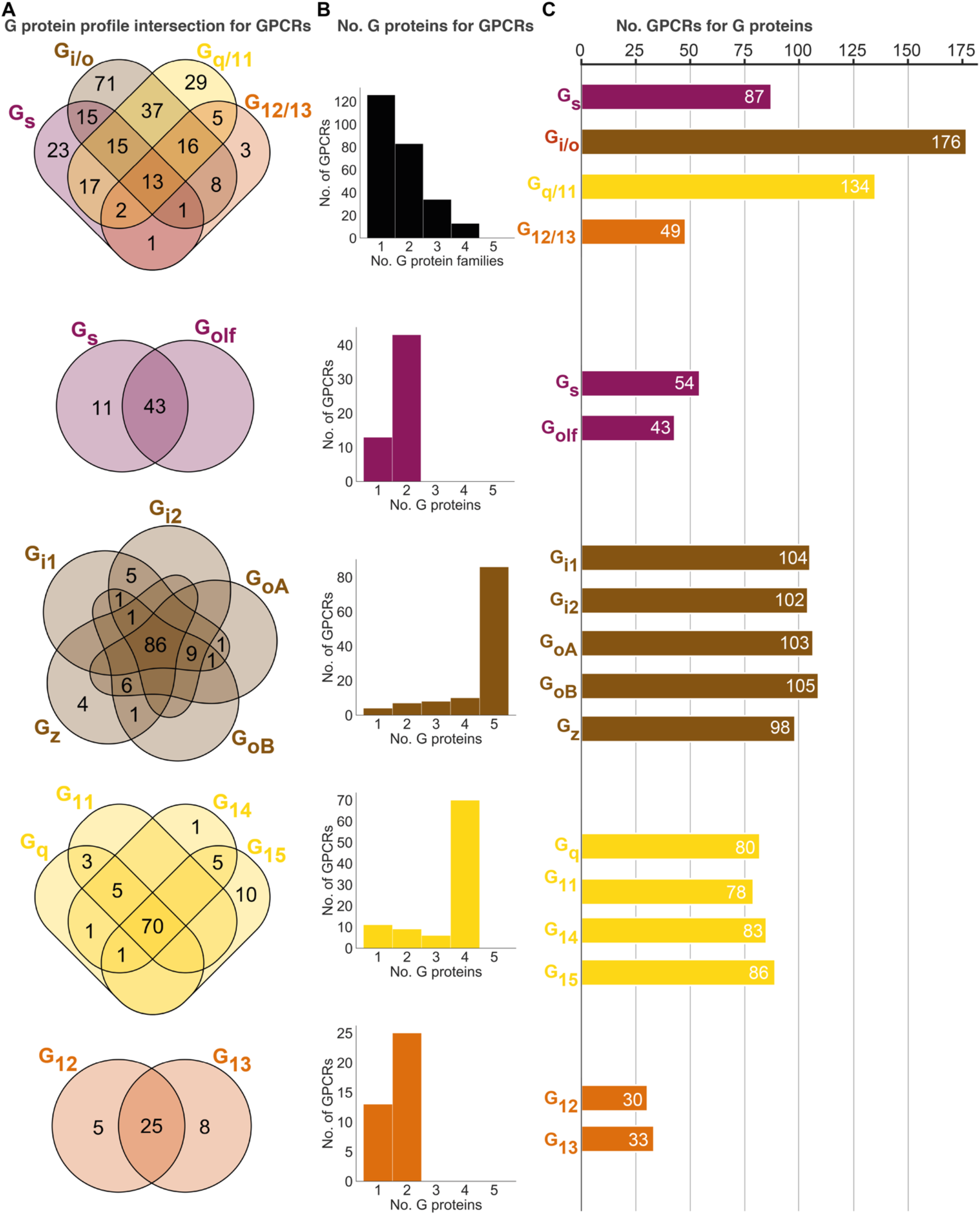
Unifying the three datasets reveals a large diversity in GPCR coupling selectivity. (**A**) GPCR-G protein selectivity on the G protein family (top) and subtype levels: Venn diagrams showing the numbers of shared and unique receptors. 0 values (no receptors) are omitted for clarity. (**B**) Receptor coupling promiscuity: Number of receptors that couple to 1 to 4 G protein families (top) and 1 to 5 subtypes. (**C**) G protein coupling promiscuity: Number of receptors that couple to each G protein family (top) or subtype. (**A-C**) Panels **B-C** are based on the couplings from the Bouvier and Inoue groups that are also supported by a second dataset and panel **A** additionally includes GtP couplings. This analysis of the G_s_ family leaves out 11 receptors tested for coupling to G_s_ but not to G_olf_, and G_olf_ couplings are only counted if there is a supported G_s_ coupling. All source data are available in tab ‘Fig_4’ in Spreadsheet S5.

Interestingly, all fully promiscuous receptors are class A GPCRs: adenosine A_1_, adrenergic α_1A,2A_ and β_1_, bradykinin B_2_, cannabinoid CB_1_, cholecystokinin CCK_1_, endothelin ET_A_, prostanoid FP, GPR4, histamine H_1_, lysophospholipid LPA_4_ and orexin OX_2_ receptors (216 class A, 11 class B1 and 5 class C GPCRs have been profiled, so far). Conversely, a G protein family has supported couplings to on average 112 GPCRs (28% of all receptors) distributed as G_s_: 87, G_i/o_ 176, G_q/11_: 134 and G_12/13_: 49 receptors (34%, 69%, 52% and 19%, respectively of all receptors, Fig. 4C). Given that 101 of the GPCR-G protein family pairs tested by the Bouvier and Inoue groups represent novel couplings (above) more couplings are expected to be identified as expanding and confirmatory studies emerge. Hence, whereas the results described here represent the currently known supported couplings, the total ‘couplome’ will undoubtedly comprise additional yet undetected and unconfirmed couplings, especially among receptors never profiled with a pan-G protein platform.

### Three-quarters of GPCR activate all G proteins belonging to the same family

Within each G protein family (rows 2-5 in Fig. 4), on average 73% of GPCRs promiscuously activate all its members (G_q/11_: 73%, G_12/13_: 66%, G_i/o_: 75% and G_s_: 80%). In contrast, activation of only one subtype of a G protein family is only observed for 11 G_s_, 4 G_z_, 1 G_14_, 10 G_15_, 5 G_12_ and 10 G_13_-coupled receptors (Fig. 2 or Spreadsheet S5). Most other receptors activate a subset of G proteins in each family. Strikingly, P2Y_1_,_4_ (G_15_), P2Y_14_ (G_z_) and GPR55 (G_13_) are fully selective also when considering G proteins from all families i.e., they only couple to a single of the 16 human G proteins (1.4*SD cut-off applied, 6 G_s_-coupling receptors are left out, as G_olf_ has so far only been tested by the Inoue group). However, the three purinergic receptors have additional couplings although not supported by a second dataset (P2Y_4_ and P2Y_14_) or above the 1.4*SD cut-off (P2Y_1_ was below) and it is possible that as the characterizations expand even fewer, or no receptors are found to engage only a single G protein.

The abundant coupling of G_i/o_- and G_q/11_ to many receptors with dual (or more) pathways suggests that these G proteins often have more versatile functions. In all, our meta-analysis of GPCR-G protein selectivity point to intriguing differences where some receptors selectively signal via a single effector whereas other GPCRs promiscuously activate all four G protein pathways. Such selective or combined profiles, in interplay with differential spatiotemporal expression (3), can be critical to achieve a specific physiological effect.

### G protein co-coupling reflects phylogeny, but all G protein families have an odd member

To assess whether the evolutionary classification dividing G proteins into four families is also representative of their pharmacology, we investigated the correlated coupling of G proteins to receptors. The overall correlation was assessed with Pearson standard correlation coefficients (Fig. 5A–B), and broken down into shared coupling/non-coupling (from Jaccard indices, Fig. 5C) and activation levels (mean log(Emax/EC_50_, Fig. 5D). We find that all three comparisons show the strongest correlations for G proteins belonging to the same G protein family (boxed in Fig. 5A, C–D). This demonstrates that the pharmacological relationships, i.e., the coupling selectivity and activation level, do indeed reflect the phylogenetic relationships. This is important, as this is how the human G proteins into four families have been classified traditionally, but it had not been tested if this classification applies to pharmacological profiles of the scales analyzed herein and likely increasingly more common in future studies.

**Fig. 5.**
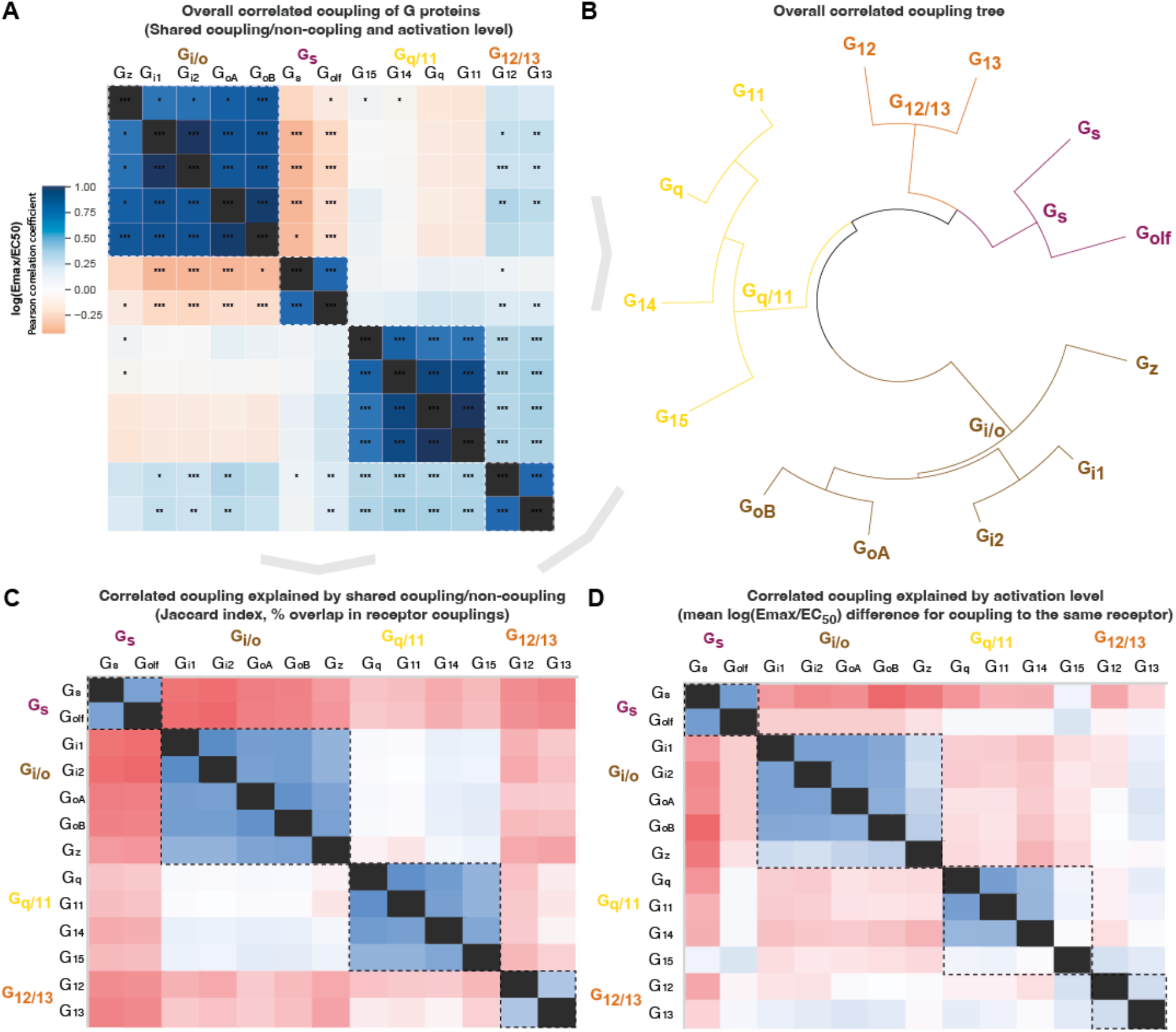
Correlated coupling of G proteins based on their receptor profiles. (**A**) Overall correlated coupling of G proteins quantified by the Pearson standard correlation coefficient, which gives a measure of the strength of the linear relationship between two G proteins (the log(E_max_/EC_50_) value of non-coupling datapoints was set to 0). Statistically significant pairwise correlations are indicated in cells by *p ≤ 0.05; **p ≤ 0.005 and ***p ≤ 0.0005. (**B**) Overall coupling correlation of G proteins shown as a tree. (**C**) Correlated coupling explained by shared coupling/non-coupling quantified as Jaccard indices (% of couplings to the same GPCRs). **(D)** Correlated coupling explained by activation level quantified as the differences in average log(E_max_/EC_50_) when a G protein pair couples to the same receptor. The values are averages of the log(E_max_/EC_50_) averages of Bouvier and Inoue values, except where data is only available in one dataset i.e., Bouvier only: G_i1_-G_i2_, G_oA_-G_oB_, G_q_-G_11_, and Inoue only: G_olf_-all G proteins. (**A, C-D**) All G protein couplings are for class A GPCRs and supported by two datasets (Bouvier or Inoue group or GtP), and their source values are available in tab ‘Fig_5’ in Spreadsheet S5. (**A-D**) For G proteins (all but G_olf_) and receptors tested by both the Bouvier and Inoue groups, an average of averages from the two groups is used. The G_oA_ and G_oB_ couplings from Bouvier were compared to the G_o_ values from Inoue which does not distinguish isoforms.

Several G protein pairs stand out with exceptionally high overall correlated coupling (Pearson standard correlation coefficient): G_i1_-G_i2_ (0.99), G_oA_-G_oB_ (0.96) and G_q_-G_11_ (0.99). Interestingly, the correlated coupling is lower for the only pairs within the G_s_ and G_12/13_ families, G_s_-G_olf_ (0.77) and G_12_-G_13_ (0.79), respectively. Furthermore, both G_i/o_ and G_q/11_ have an ‘odd’ member, G_z_ and G_15_, respectively (mean of 0.78 and 0.76 to other family members, respectively). The fact that all G protein families have an odd member shows that all signaling pathways have a transducer toolbox allowing them to differentiate signaling.

### G_s_ or G_i/o_ coupling is selective while G_12/13_ is promiscuously activated with G_i/o_ and G_q/11_

The G_s_ and G_i/o_ families have an inverse correlation in all of the overall, coupling/non-coupling and activation level comparisons (darkest red in Fig. 5A, C–D). This means that G_s_ and G_i/o_ rarely co-couple to GPCRs and, when they do, they do so with a large difference in their strength of activation. This inverse correlation where only one or the other G protein is activated is in agreement with their function, as G_s_ stimulates, and G_i/o_ inhibits production of the same cellular second messenger, cAMP. This allows opposite physiological responses to be mediated with temporal selectivity in the same cell by activating different receptors at different times. Differential engagement of the G_i/o_ family could also have functional consequences through other pathways, as G_i_ was recently shown to be required for scaffolding β-arrestin binding and signaling for some receptors (9).

For G_12/13_, we instead find a positive correlation with the G_i/o_ and G_q/11_ families. This is in agreement with Fig. 4 (first Venn) which shows that only 3 GPCRs couple to only G_12/13_ and 3 couple to the rare G_12/13_- G_s_ family pair, whereas the vast majority of GPCRs, 38 couple to G_12/13_ also couple to the G_i/o_ and/or G_q/11_ family. Furthermore, we find an intriguing unique selectivity mechanism for the G_i/o_ and G_q/11_ families. These families have the most frequent co-coupling (light blue in Fig. 5C) but have a high difference in average activation levels (red in Fig. 5D). Together, this shows that selective binding of only G_i/o_ or G_q/11_ is uncommon, but selectivity can be achieved by differential levels of activation. This possibility to achieve selectivity via differential activation is likely important, as many GPCRs, 81 couple to both the G_i/o_ and G_q/11_ families. Their high difference in activation levels allow for a selective activation, which could shift upon the presentation of alternative endogenous or surrogate ligands with a signaling bias. Furthermore, it should be pointed out that many reported dual couplings to G_i_ and G_q_ may be because PLC, a direct effector of G_q_, can also be activated by G_βγ_ from G_i_ (i.e. sensitive to the G_i_ inhibitor pertussis toxin). Thus, the concept of potential scaffolding of PLC by G_q_ may be required for G_i_-derived G_βγ_ activation (10).

### Differential tissue expression gives G proteins in the same family large spatial selectivity

To study how tissue expression may influence G protein selectivity, we analyzed consensus transcript expression levels for 50 tissues and 16 organs (here aggregated into 8) from the Human Protein Atlas (11), which also includes data from the Genotype-Tissue Expression (GTeX) project (12) (Fig. 6). We find that the most ubiquitously expressed G proteins are G_s_ and G_i2_ which are expressed in all 50 tissues and all organ categories at a level that is within the first quartile of all normalized transcripts per million (nTPM) values. In contrast, G_i3_, G_t1_, G_t2_, G_t3_, and G_14_ have none or only one tissue with a first quartile expression. To facilitate comparison across G proteins, we employed HPA’s normalized transcripts per million (nTPM) values after z-score transformation of each G protein across tissues. A z-transformation of the same G proteins visualizing the relative tissue levels of expression, shows a very focused expression of the transducins, G_t1_ and G_t2_, in the retina and gustducin G_t3_ in the gastrointestinal tract reflecting their role in vision and taste, respectively (Fig. S2A). Notably, each G protein family has at least one subtype that is preferentially expressed in the brain: G_s_: G_olf_, G_i/o_: G_o_ and G_z_, G_q/11_: G_q_ and G_12/13_: G_12_ (Fig. 6 and Fig. S2A).

**Fig. 6.**
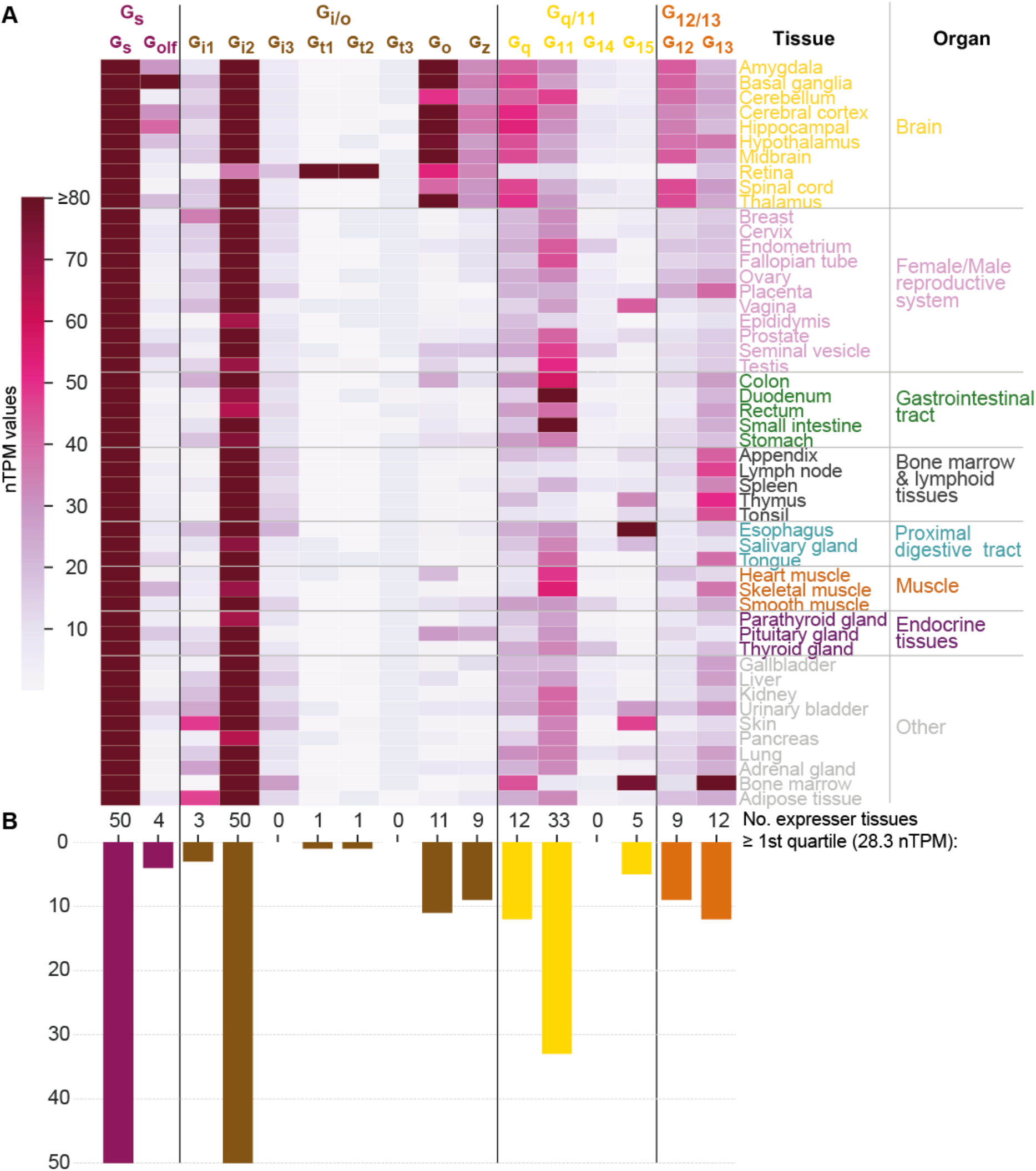
G protein tissue and organ expression profiles. (**A)** Expression heatmap of all 16 human G proteins across 50 tissues and 16 organs (here grouped in 8 categories) extracted from the Human Protein Atlas (11), which also includes data from the Genotype-Tissue Expression (GTeX) project (12). The coloring denotes the normalized transcripts per million (nTPM) for each G protein and tissue capped at the median of all maximum G protein expression values (G_t3_ at 80 nTPM in retina). (**B)** Number of tissues for which the given G protein has an expression the first quartile expression threshold (28.4 nTPM) across all 16 G proteins and 50 tissues.

Given that the tissue expression varies largely for G proteins, we next sought to determine to what extent their tissue expression correlations differ from the traditional four G protein families that represent phylogeny and GPCR coupling (Fig. 5) relationships. To this end, we calculated the co-expression for each G protein pair using Pearson standard correlation coefficients (Fig. S2B). Notably, we find that all four G protein families have members that group apart in the dendrogram (the two members of the G_s_ family, G_s_ and G_olf_, do not cluster adjacently but not as a pair). Instead, we find a coherent cluster for the G proteins that are predominantly expressed in the brain, G_olf_, G_o_, G_z_, G_q_ and G_12_ and cover all four families. Furthermore, among the remaining co-expressing G proteins, three out of the four closest pairs also span families (G_i3_-G_15_, G_i2_-G_13_ and G_t3_-G_11_ but not G_t1_-G_t2_). Taken together, this shows that there is no overall clustering of subtypes within the G protein families, which instead exploit differential expression to gain spatial selectivity. For example, in the G_i/o_ family G_o_ and G_z_ are restricted to brain regions, while the other subtypes have four different peripheral profiles.

## Technological considerations – Biosensor sensitivity

### Bouvier’s and Inoue’s biosensors appear more sensitive for G_15_ and, G_s_ and G_12_, respectively

To evaluate the sensitivity of the Bouvier and Inoue biosensors for different G proteins, we compared their observed couplings for the 70 common receptors to identify ‘unique’ and ‘missing’ couplings (defined in Table 1). For G_15_, Bouvier reported 30% unique couplings, several of which were validated in Ca_2+_ assays (3), whereas Inoue instead misses 13% of couplings present in GtP (Fig. 7). This indicates that G_15_ couplings are underrepresented in Inoue’s data and in literature (annotated in GtP). The few published couplings for G_15_ is likely explained by its lack of expression in the cells most used for *in vitro* experiments, such as HEK293 cells (3), the lack until recently of enough sensitive sensors to directly measuring G_15_ activity, and by the lack or weak (at very high concentration) effect on this subtype by G_q/11_ inhibitor tools YM-254890 and FR900359, respectively (13).

**Fig. 7.**
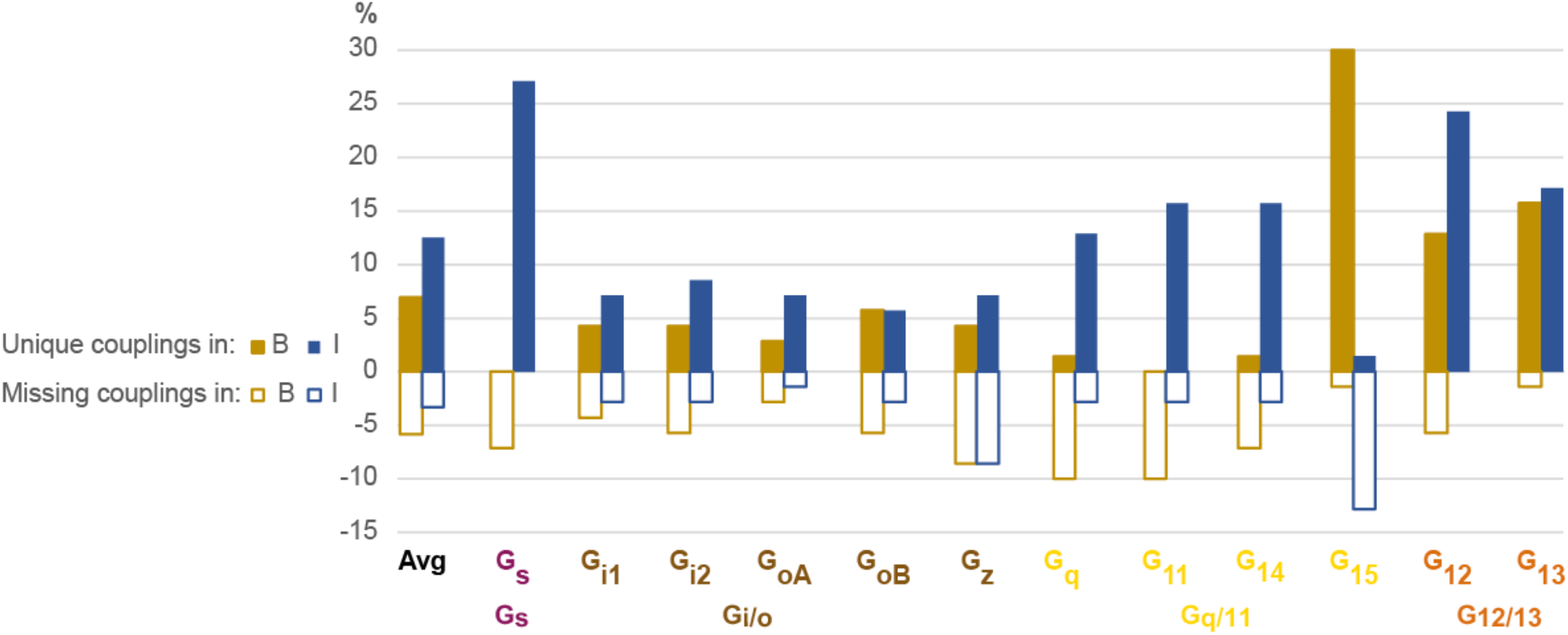
Unique and missing couplings in the Bouvier and Inoue datasets. Unique and missing couplings (defined in Table 1) among all 70 receptors common to Bouvier, Inoue and GtP (the 100% includes non-coupling receptors). Missing couplings are shown as negative values. For GtP, we consider coupling to a G protein subtype possible if a coupling has been observed for the respective family. Unique couplings are hidden by default in the online G protein couplings browser in GproteinDb, as they await the independent support by a second group (5).

For G_s_ and G_12_, 27% and 24%, respectively of Inoue’s couplings are unique, whereas Bouvier instead misses 7% of G_s_ and 6% of G_12_ couplings in GtP. These two G proteins have smaller assay windows which may explain their lower representation in the Bouvier dataset (tab ‘DataStats’ in Spreadsheet S1). Furthermore, for G_q_, G_11_ and G_14_, 13%, 16% and 16%, respectively of Inoue’s couplings unique. In contrast, Bouvier is missing 10%, 10% and 7%, respectively of the known couplings in GtP which indicates an underrepresentation or alternatively that literature has mistaken G_i_-derived βγ activation of the PLC-IP3-Calcium signaling pathway for G_q_ mediated signaling (14–17). Notably, although Inoue used the wildtype protein for G_q_, it has a similar or higher frequency of unique couplings compared to all other measured G proteins (G_q_: 13% vs. G_i1_: 7%, G_i2_: 9%, G_oA_: 7%, G_oB_: 6%, G_z_: 13%, G_11_: 16%, G_14_: 16%, G_15_: and G_13_: 17%), except G_s_ and G_12_. This suggests that the many unique couplings in Inoue’s dataset may be due to high sensitivity rather than the use of chimeric G proteins carrying over of G_q_ coupling to the other G proteins.

To investigate how unique couplings are affected by weak couplings and biosensor sensitivity, we compared their efficacies and potencies (rows 106-110 in tab ‘BIG-QualComp’ in Spreadsheet S3). We find that the average E_max_ values are 18% and 15% lower for unique than for supported couplings in the Bouvier and Inoue datasets, respectively. Furthermore, their average pEC_50_ values are 0.4 and 1.5 log units – 2.5-fold and 32-fold, respectively – lower than the supported couplings. This indicates that a part of the differences across datasets is due to weak couplings that can only be detected in the most sensitive assays for the given G proteins and are difficult to distinguish from basal levels (this study used a cut-off of E_max_>1.4 SDs over basal response). However, couplings may also be overrepresented and determining couplings definitely would require further studies employing independent (‘orthogonal’) high-sensitivity biosensors. Until then, we recommend the requirement of two independent studies to support of a GPCR-G protein coupling, as is the criterion used herein for the novel couplings and the default in the online G protein couplings resource (1).

## Discussion

Given that researchers are now faced with three large G protein coupling datasets varying in coverage and couplings, we established a common normalization protocol making quantified values comparable across data sources and a common coupling map. Previous analyses of GPCR-G protein selectivity have all been based on a single dataset from the GtP (18) database or the Inoue (2) or Bouvier (3) laboratories, whereas the analysis herein combined these datasets to establish couplings supported by two studies (including GtP annotations of multiple literature reports) for a total of 256 GPCRs. This provides a sizeable and reliable reference dataset of supported quantitative couplings suitable for any study.

By analyzing the common coupling map, we find that 101 novel couplings are supported by both the Bouvier and Inoue datasets but were not known in GtP which annotates coupling from the literature. We also find that very few receptors, 13 (5%) promiscuously couple to all four G protein families whereas the receptor numbers instead increase several-fold along with selectivity: 34 triple-, 83 double- and 126 single-family coupling receptors (Fig. 4B). As more large coupling datasets become available, this is likely to shift somewhat from less selectivity towards higher promiscuity. Furthermore, the finding that on average only 27% of GPCRs activate only a subset of the G proteins in each family (Fig. 4) opens the possibility that signaling bias on the level of specific subtypes may be much more frequent than currently appreciated. G proteins that belong to the same family can have diverse functional outcomes pertaining to effector engagement selectivity and kinetic profiles (19, 20), suggesting that bias within a G protein family may have physiological and/or therapeutic implications (21). The observation that many GPCRs can couple to more than one family of G proteins but still can show selectivity between members of a same family, opens important questions concerning the structural determinants of such multifaceted selectivity profiles.

The GPCR-G protein couplings analyzed herein represent the overall couplings that are possible, but may differ largely physiologically across specific tissues. Our analysis showed a tremendous span in how many tissues that express a given G protein. Some G proteins, including the transducins (G_t1_ and G_t2_) and gustducin (G_t3_) have a restricted expression in mainly one tissue (retina and the gastrointestinal tract, respectively), whereas the most ubiquitously expressed G proteins, G_s_ and G_i2_ are highly expressed in all 50 analyzed tissues. Notably, G proteins belonging to the same family differ largely in their tissue profiles for example, each G protein family has a single or two subtypes that is/are preferentially expressed in the brain. Given that the activity (here log(E_max_/EC_50_) values) can vary substantially across G protein family subtypes (Fig. 2), this provides cells with a mechanism to shift the system bias towards different intracellular signaling pathways. Such system bias can be modulated further through differential expression of also the receptors and multiple downstream intracellular effectors (22). A recent study found a strong spatial correlation of summed mRNA expression levels for G_s_-, G_i/o_-, and G_q/11_-linked receptors in humans, macaques, and mice (23). This suggests that the expression patterns of receptors linked to these major G protein families are strongly interdependent and could therefore act together to balance each other in specific tissues. Future studies would be needed to profile the couplings of natively expressed GPCRs and G proteins across different cell types. Meanwhile, a strategy could be to analyze GPCR-G protein selectivity for a subset of G proteins that are selected based on the highest expression in the tissue of interest. For example, analysis of G_i/o_-mediated signaling in the brain could focus on G_o_ and G_z_ based on their high expression, and the fact that G_o_ and G_z_ have quicker and slower nucleotide exchange, respectively than other member of this family. Such separate meta-analysis of G proteins may be facilitated by the filtering options in the coupling maps in Spreadsheet S5 and online in the G protein couplings browser in GproteinDb (5).

The Inoue and Bouvier datasets represent the first steps to systematically characterize large G protein coupling profiles. Other laboratories may use other biosensors to expand the coverage of the coupling map, and our study facilitates this by indicating which of the Bouvier and Inoue biosensors that have the highest sensitivity for each G protein (Fig. 7). However, future studies should also employ different biosensors and systems, as independent support is critical for validation and for distinguishing novel couplings from false positives among the many unique couplings (Fig. 7 and Spreadsheet S3). Of note, several biosensors have recently been published (7, 8, 24) (Gαβγ sensors first described in (25–30)). Whereas a detailed comparison is out of the scope of this study, their pros and cons have recently been reviewed (31) and researchers should strive to use the most native proteins and expression, physiologically relevant tissues/cells (32) and assays with sufficient sensitivity and window. Of note, most studies annotated in GtP have used downstream measurements, which are amplified, and may therefore differ from those obtained by biosensors that measure G protein activation (further discussed in (3)).

In all, our cross-dataset analysis has established a protocol and reference set aiding GPCR-G protein coupling studies. The selectivity profiles are the most comprehensive to date spanning 256 receptors with a very diverse activation of a single to all G protein pathways, and presented in a dedicated online G protein couplings browser in GproteinDb (18). The analyses and data presented herein will be very valuable to illuminate undercharacterised pharmacological phenomena such as constitutive activity (33), pre-coupling of G proteins (26, 34) and ligand-dependent biased G protein signaling (35), and to uncover their underlying determinants. They also present the foundation to integrate more coupling data as future studies expand the characterization of the ‘couplome’.

## Methods

### Study design

The primary objective of this study was to generate a unified map of G protein couplings (Fig. 2) across the three available large datasets from the Bouvier (3) and Inoue (2) groups, and the Guide to Pharmacology database (1), respectively. To do this, we identified the Emax standard deviation cut-off, quantitative normalization protocol and aggregation of G proteins into families giving the best possible agreement between the GPCR-G protein couplings from the Bouvier and Inoue groups. This analysis also involved assessing the agreement of the Bouvier and Inoue group datasets and determining the number of high-confidence novel couplings supported by these two datasets but not reported in the Guide to Pharmacology database. Furthermore, it included a benchmarking of techniques to determine which biosensor produces the most reproducible qualitative coupling determination (coupling vs. non-coupling) for each G protein family. To this end, we compared the three datasets to pinpoint, for each dataset and G protein family, the fraction of couplings supported by another dataset. The scientific analyses based on the map feature the most comprehensive analysis to date of GPCR-G protein selectivity. This was done by intersecting the 11 G proteins tested by both Bouvier and Inoue, within and across their families, with respect to the common and unique receptors that they couple to. Finally, the receptor profiles were also used to classify G proteins and to determine any co-correlation between different G protein subtypes and families to GPCRs.

### Coupling datasets

Updated spreadsheets containing the pEC_50_, E_max_, basal signals and standard deviation (SD) values were supplied by Asuka Inoue. Basal signals, spontaneous AP-TGF-α elease (in % of total AP- TGF-α expression) were recalculated from raw data of the previous coupling-profiling campaign (2) and their SD values were computed from independent experiments (n ≥ 3). This file contains the neurotensin 1 (NTS_1_) and Thyrotropin-releasing hormone (TRH_1_) receptors not included in the previous publication (2). In the Inoue dataset, the protease-activated receptors PAR3-4 had negative pEC_50_ values (concentration of mU/ml because the ligand, thrombin, was supplied with its enzymatic activity). For the easiness of integration into the coupling map, we added a value of 10 to their pE_C50_ values. Data qualities (sigmoidal curves) for the individual GPCR-G protein pairs were manually inspected and concentration-response curves that did not converge nor exceed a threshold (typically, 3% AP-TGF-α release) were regarded as no activity.

The Bouvier dataset (n ≥ 3) contained some datapoints that were included or excluded based on dedicated analyses. Firstly, we excluded ligand-promoted responses of overexpressed receptors that were equivalent to those of endogenously expressed receptors (yellow fill in tab ‘B-EmEC’ in Spreadsheet S1). Secondly, couplings with only approximate E_max_ and pEC_50_ values because the dose-response curve did not converge were only included if supported, and not contradicted, by the Inoue and/or GtP datasets (orange fill in tab ‘B-EmEC’ and analyzed separately in tab ‘UnconvergedDRV’ in Spreadsheet S1). This is because a coupling with an unprecise quantitative value is better than no coupling, especially when making qualitative comparisons (coupling vs. non-coupling). Based on these criteria seven couplings were included: 5-HT_1D_-G_z_, BLT_1_-G_11_, FFA3-G_13_, GPR_4_-Gi_1-2_, GPR84-G_oA_ and κ-G_12_, and six couplings were excluded: CCR5-G_14_, CXCR5-G_q_/G_11_/_G14_, GPR183-G_13_ and κ-G_13_.

### Standard deviation cut-off

We made a special investigation of couplings that have a full dose-response curve but an E_max_ less than 2 standard deviations (SDs) from the basal signal (red fill in tab ‘B-EmEC’ and analyzed separately in tab tabs ‘B-<2SDs’ and ‘I-<2SDs’ in Spreadsheet S1). To achieve the best possible separation of putative false and real but weak couplings, we identified the threshold value (a number of SDs from basal signal) that gives the best agreement between the Bouvier and Inoue dataset among the common receptors and G proteins tested by both groups. The obtained cut-off, 1.4 SDs was applied as a filter to exclude all Bouvier and Inoue E_max_ and pEC_50_ values for couplings below this cut-off (columns with ‘>1.4SD’ in the heading in tabs ‘B-EmEC’ and ‘I-EmEC’ in Spreadsheet S1). As a note, while intra-day measurement error is small for the TGF-α shedding assay (typically, 1-2% AP-TGF-α release), inter-day variability varies widely depending on cell conditions. Since the SD represents inter-day variability, the basal SD cut-off removes more couplings than the SD cut-off of ligand-induced signal or the significance of individual experiments. As a consequence, some of manually annotated couplings in the Inoue dataset are regarded as non-coupling by the basal SD cut-off criteria, including for P2RY_2_ and P2RY_6_ that had no couplings above this cut-off.

### Generating a subset of comparable GPCR-G protein couplings

To enable qualitative comparison of corresponding datapoints in the datasets from the Bouvier (3) and Inoue (2) groups, we used the subset of 70 GPCRs present in both datasets and belonging to the same class, A (removed only two receptors from class B1). The quantitative comparisons focused on a smaller subset of 51 GPCRs tested with the same ligand and excluded non-coupling GPCR-G protein pairs, as they could not be represented by 0 values (due to e.g., an underrepresentation of couplings in a given datasets and G proteins, see Results). All analyses herein included the 12 G proteins: G_s_, G_i1_, G_i2_, G_oA_, G_oB_, G_z_, G_q_, G_11_, G_14_, G_15_, G_12_ and G_13_. G_olf_ and G_i3_ could not be analyzed, as they had not been tested by the Bouvier group (Table S1). The Inoue data for the pairs G_i1-2_, G_oA-B_ and G_q_ and G_11_, respectively, were generated with identical chimera inserting the Gα C-terminal hexamer into a G_q_ backbone (2). Qualitative analyses compared the presence or absence of each GPCR-G protein coupling while the quantitative analyses were limited to common G protein couplings, i.e., data points in which both the Bouvier and Inoue groups generated a pEC_50_ and E_max_ value.

### Validation of comparability when E_max_ or reference agonist differ

To get an overview of the distribution of data in the two datasets, we determined the pEC_50_ and E_max_ mean, median, min, max, span and standard deviation values and plotted box and whiskers plots for each G protein (Spreadsheet S1). This showed that the E_max_ values vary much more across the different G proteins than pEC_50_ values. The E_max_ variation is largest in the Bouvier data wherein G_15_ ranges across three orders of magnitude (16 to 1,067). The Bouvier dataset has low means and spans for G_s_ (42 and 47) and G12 (71 and 101) indicating a narrow assay signal window (low signal-to-noise). We find that minimum-maximum normalization (to 100%) of each G protein across receptors gives a more uniform distribution (SDs for Bouvier: 19-31 and Inoue: 22-26, rightmost plot pair in Spreadsheet S1).

Whereas both studies tested a majority of receptors with their endogenous ligand, surrogate agonists were used for 15 and 4 GPCRs in the data from the Inoue and Bouvier groups, respectively. Although those ligands were selected for their reference character with similar pharmacology to the endogenous ligand, they could introduce differences in a receptor’s G protein profile due to ligand-dependent signaling bias. Analysis of GPCR-G protein couplers and non-couplers (tab ‘BI’ in Spreadsheet S2) shows that on average 74% and 71% agreeing qualitative couplings (i.e., coupling vs. non-coupling GPCR-G protein pairs) for receptors tested with the same and different agonists, respectively. This is a rather small difference providing confirmation that receptors tested with different ligands can be compared on the qualitative coupling/non-coupling level and be included in the comparison of the different G protein coupling datasets and determination of novel G protein couplings.

### Dataset integration into a unified coupling map – Normalization and log(E_max_/E_C50_) values

To enable quantitative correlation of the Bouvier and Inoue couplings, we further filtered the 70 common GPCRs to yield 51 common class A GPCRs tested with the same ligand (excluding 29 receptors tested with different ligands and two class B1 GPCRs). To assess the value of normalization, we calculated the average ‘similarity’ (Bouvier/Inoue ratio) of individual values (tabs ending with ‘-sim’ in Spreadsheet S4) and the ‘linear correlation’ (r2 value) of each receptor across all G proteins (reported below as averages of individual couplings and G proteins, respectively) (tabs ending with ‘-r2’ in Spreadsheet S4). Linear correlation was only done for receptors with at least three common G proteins/families. Minimum-maximum normalized E_max_ values were represented as percentage values while decimal values (0-1) were used for the calculation of log(E_max_/E_C50_) values as recommended in (4). For both the Bouvier and Inoue groups, the minimum and maximum represent the signal without (0%) and with (100%) an agonist, respectively, in each experiment replica. The minimum E_max_ value was therefore set 0 while the maximum was set to the highest value for the given G protein (first, use normalization) or receptor (second, tested but not used normalization).

Minimum-maximum normalization of E_max_ measurements increased the average from 0.22 to 0.61 and the linear correlation r^2^ value from 0.28 to 0.30. In contrast, non-normalized pEC_50_ measurements have the most similar values (0.89 compared to 0.66) and an identical linear correlation (0.37). We combined the minimum-maximum normalized E_max_ and non-normalized EC_50_ values into log(E_max_/EC_50_) (4). The log(E_max_/EC_50_) values have an average similarity that is nearly as high (0.86 compared to 0.89) and an average linear correlation that is better (0.41 compared to 0.37) than for pEC_50_, the best individual measure. We did not apply a double normalization, i.e., also across G proteins after across receptors, as this worsens the E_max_, pEC_50_ and log(E_max_/EC_50_) value similarity (from 0.61 to 0.39, 0.89 to 0.42 and 0.86 to 0.47, respectively) and linear correlation of each G protein across receptors (from 0.11 to 0.05, 0.49 to 0.20 and 0.46 to 0.23, respectively.

### Dataset integration into a unified coupling map – G protein family aggregation

Given that G proteins belong to families that are functionally grouped by sharing downstream signaling pathways, we next investigated the best member-to-family aggregation scheme – specifically, whether the most comparable G protein family values are obtained if using the maximum value from any single subtype or the mean of all subtype members. We found that aggregation using max rather than mean values gives a better similarity for E_max_ (0.69 vs. 0.65), marginally lower similarities for pEC_50_ (0.89 relative 0.90) and identical similarities for log(E_max_/EC_50_) (0.88). However, max performs better overall than mean in the comparisons correlating a receptor across G proteins; E_max_ (0.63 and 0.53), pEC_50_ (0.56 and 0.56) and log(E_max_/EC_50_) (0.65 and 0.59) or one G protein across receptors; E_max_ (0.29 and 0.20), pEC_50_ (0.71 and 0.74) and log(E_max_/EC_50_) (0.69 and 0.71). Notably, the correlation of each receptor across max-aggregated G protein families compared to non-aggregated subtypes is much stronger when considering any of the three pharmacological parameters: E_max_ (0.63 vs. 0.36), pEC_50_ (0.56 vs. 0.30) and log(E_max_/EC_50_) (0.65 vs. 0.41). Altogether, this establishes the highest value (max) as the aggregation that yields the most comparable quantitative value for G protein families.

### Tissue expression analysis

Consensus transcript expression levels were extracted for 50 different tissues aggregated into 16 organs or systems in the Human Protein Atlas (HPA) (11), which also includes data from the Genotype-Tissue Expression (GTEx) project (12). The G protein expression was quantified in the form of normalized transcripts per million (nTPM), which is a consensus normalized transcript expression value (see https://www.proteinatlas.org/about/assays+annotation#rna). The correlation analysis was performed a default Pearson correlation analysis using Seaborn’s clustermap.

### Statistical analysis

The aggregated sample size is n=3 or higher for all analyzed GPCR-G protein couplings. The specific sample size for each such coupling is described in the original articles reporting these data and referenced in the present manuscript. The Bouvier dataset contained some datapoints that were included or excluded based on dedicated analyses (see ‘Coupling datasets’ above). For all figures, the values for *N*, *P*, and the specific statistical test is included in the figure legend or associated manuscript text. For Fig. 5, pairwise correlation of log(Emax/EC_50_) values was assessed as a measure of the strength of the linear relationship between two pathways. The pairwise distances between pathways were calculated in python scipy package using the *spatial.distance.pdist* method using the *correlation* metric. Statistical significance was determined by Pearson standard correlation coefficient with a two-tailed p-value as implemented in scipy’s *stats.pearsonr* method.

## Acknowledgments

Nevin Lambert is acknowledged for fruitful discussions. We thank Ayumi Inoue (Tohoku University) for reanalyzing the raw data of the TGF-α shedding assay to obtain SD values for the individual GPCR-G protein pairs.

## Additional information Funding

Canada Research Chair in Signal Transduction and Molecular Pharmacology (MB); Canadian Institutes of Health Research grant FDN-148431 (MB); Lundbeck Foundation grants R218-2016-1266 and R313-2019-526 (DEG); Novo Nordisk Foundation grant NNF18OC0031226 (DEG); BINDS program grant JP20am0101095 (AI); LEAP program grant JP20gm0010004 (AI); Japan Agency for Medical Research and Development (AMED) (AI); Takeda Science Foundation (AI); Uehara Memorial Foundation (AI);

## Author contributions

Conceptualization: DEG; Methodology: DEG; Validation: ASH, DEG; Formal Analysis: AI, CA, ASH, DEG; Investigation: ASH, CA, DEG; Resources: CG; Data Curation: AM, CA, CN, DEG; Writing – Original Draft: DEG, MB; Writing – Review & Editing: AI, ASH, CA, DEG, MB; Visualization: ASH, DEG; Supervision: DEG, MB; Project Administration: DEG; Funding Acquisition: AI, DEG, MB.

## Data and materials availability

All data are available in GproteinDb (https://gproteindb.or), GitHub (https://github.com/protwis/gpcrdb_data) and Spreadsheets S1-5. No code was developed for this manuscript which instead used the existing resources of GproteinDb (5) and GPCRdb (36) resources. All open-source code can be obtained from GitHub (https://github.com/protwis/protwis) under the permissive Apache 2.0 License (https://www.apache.org/licenses/LICENSE-2.0).

## Supplementary Information

**Fig. S1.**
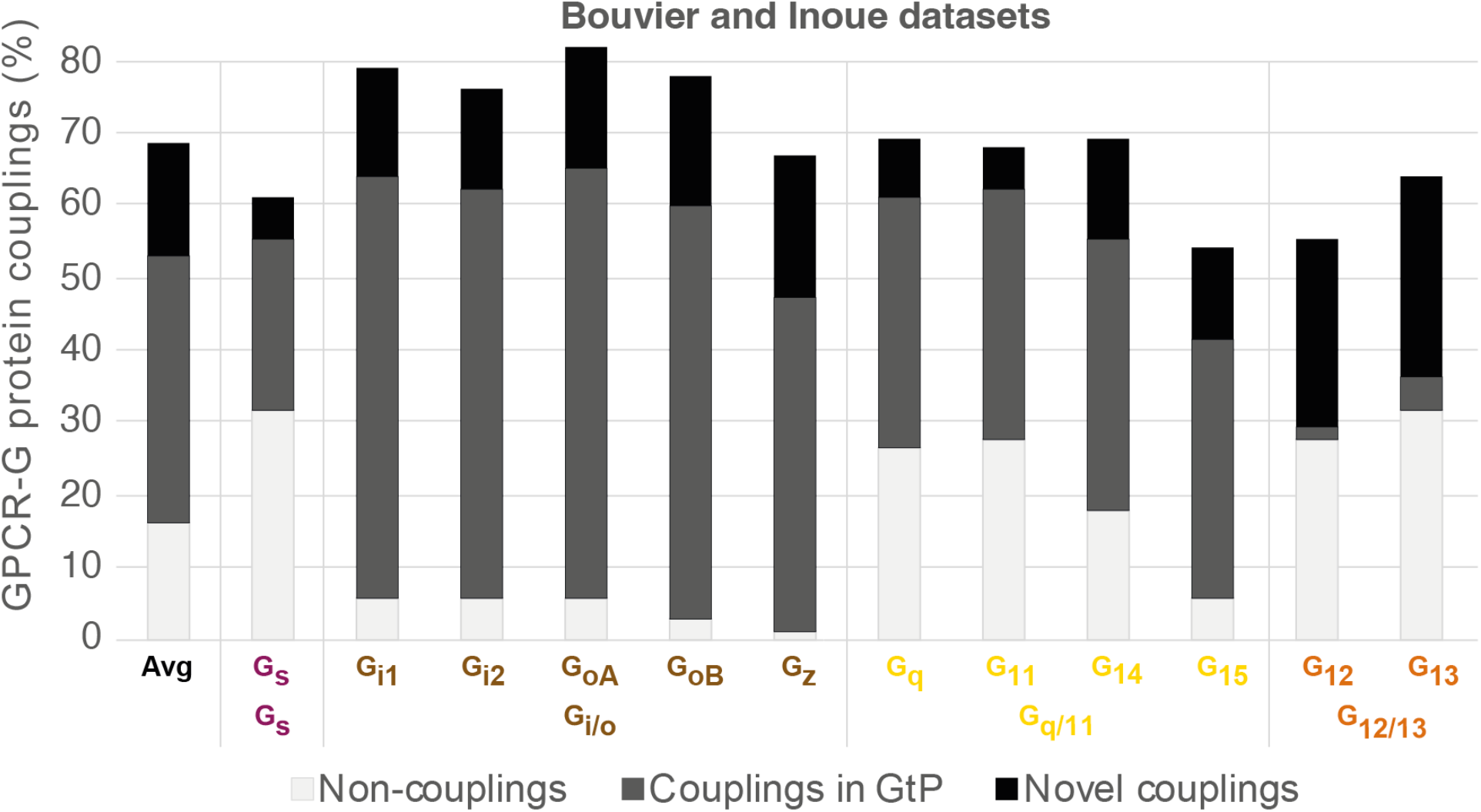
Common GPCR-G protein couplings in the Bouvier and Inoue datasets. Black bars represent couplings not in GtP and therefore considered novel (novel couplings are listed in Fig. 3). Frequencies are percent couplings among all receptors common to Bouvier and Inoue (the 100% includes non-coupling receptors). G_i3_ and G_olf_ (chimeras) are not included herein as they were only tested by the Inoue group. Inoue used the same chimeras to represent the pairs G_oA_-G_oB_, G_i1_-G_i2_ and G_q_-G_11_ (Table S1).

**Fig. S2.**
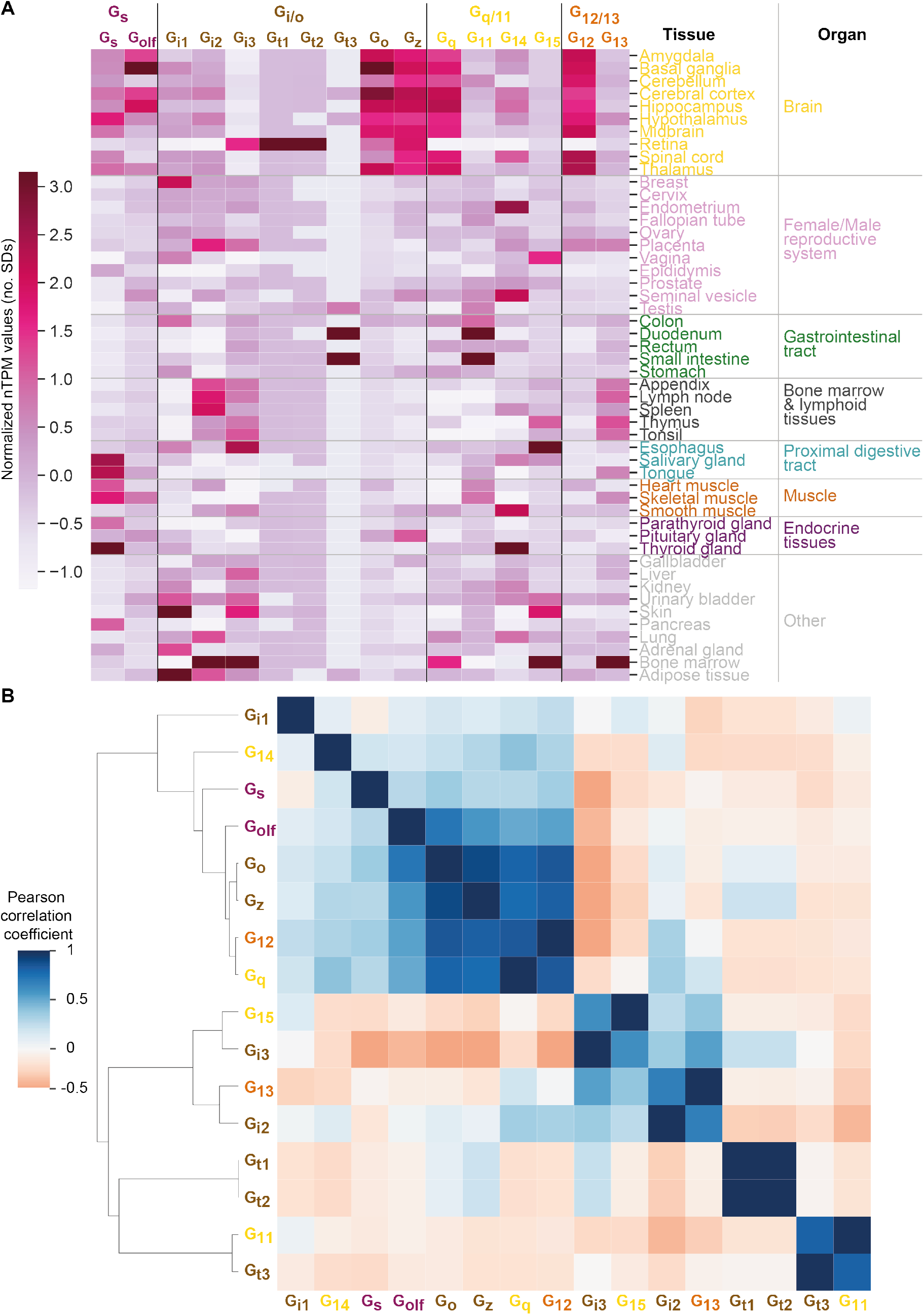
Correlated expression of G proteins. (**A)** Expression heatmap like in Fig. 6 but with a z-transformation across tissues, thereby visualizing their relative expression for each G protein. (**B)** Correlated G protein expression across tissues as a function of normalized transcripts per million (nTPM), which is a consensus normalized transcript expression value across the Genotype-Tissue Expression (GTEx) and Human Protein Atlas datasets (see Methods). Pairwise correlation of nTPM expression values by Pearson standard correlation coefficient gives positive and negative correlations between G protein pairs. Expression clusters are distinct from the G protein families, which are denoted with color-coded labels.

**Table S1.**
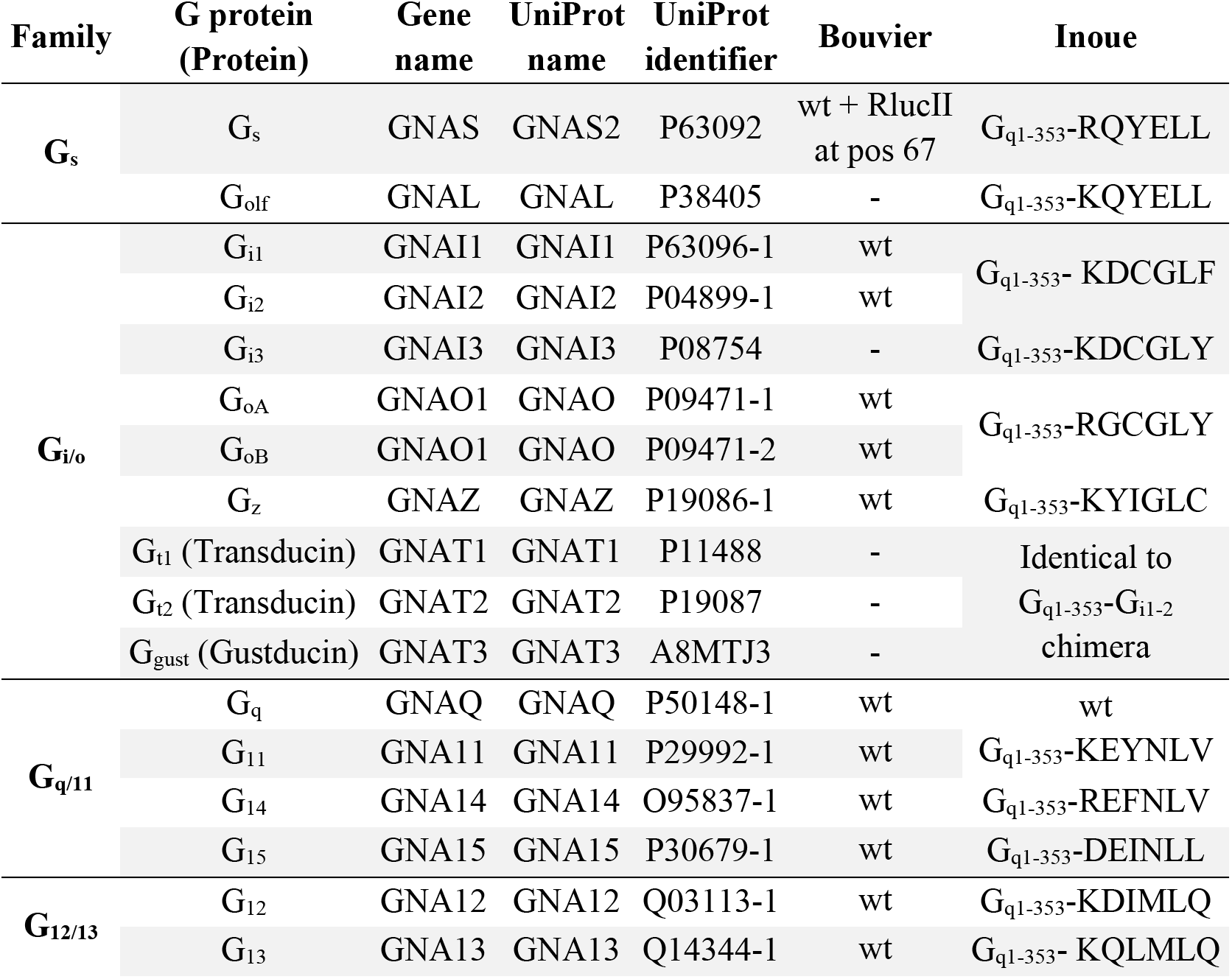
G proteins tested by Bouvier et al. and Inoue et al. The two published datasets contain 11 common G proteins (G_s_, G_i1_, G_i2_, G_o_, G_z_, G_q_, G_11_, G_14_, G_15_, G_12_ and G_13_), two Inoue et al. specific G proteins (G_olf_ and G_i3_) and two Bouvier specific isoforms (G_oA_ and G_oB_) of the Go protein that spring from the same gene (GNAO1). Inoue et al. G_q_ chimera replacing the six C-terminal amino acids have identical sequences for G_i1-2_, G_oA-B_, and G_q_ and G_11_. All analyses herein used the 11 common G proteins, a G_o_ average of the G_oA_ and G_oB_ isoforms and left out G_olf_ and G_i3_ (not present in Bouvier et al.) while identical chimeras from Inoue et al. were used to represent the both members of the pairs G_i1-2_, G_oA-B_ and G_q_ and G_11_, respectively. Abbreviations: wt = wildtype. RlucII = Renilla luciferase 2.

## Titles Of Additional Supplementary Material Files

**Spreadsheet S1. SD cut-off and pE_C50_ and E_max_ distributions.**

This spreadsheet contains a calculation of a standard deviation (SD) cut-off that gives the best agreement between G protein couplings from the Bouvier and Inoue groups. The cut-off, 1.4 represent the number of standard deviations of a G protein coupling average (at least from triplicates) from the basal signal of the given receptor. The spreadsheet also contains a DataStats tab detailing the distribution of raw and normalized pE_C50_ and E_max_ values.

**Spreadsheet S2. Assessment of effect of different ligands.**

This spreadsheet contains a comparison of the relative average difference obtained for G protein couplings when determined with and without using the same ligand in the Bouvier and Inoue groups.

**Spreadsheet S3. Qualitative coupling comparisons.**

This spreadsheet contains a qualitative comparison (coupling and non-coupling) of GPCR-G protein pairs from the Bouvier and Inoue groups, and from the Guide to Pharmacology database.

**Spreadsheet S4. Normalization and aggregation into G protein families.**

This spreadsheet contains a comparative analysis of which normalization and aggregation into G protein families that gives the best agreement between G protein couplings from the Bouvier and Inoue groups.

**Spreadsheet S5. Unified coupling map.**

This spreadsheet contains a unification of G protein couplings from the Bouvier and Inoue groups, and from the Guide to Pharmacology database into a common coupling map.

